# Cell wound repair requires the coordinated action of linear and branched actin nucleation factors

**DOI:** 10.1101/2022.04.14.488397

**Authors:** Justin Hui, Mitsutoshi Nakamura, Julien Dubrulle, Susan M. Parkhurst

## Abstract

Cells are subjected to a barrage of daily insults that often lead to its cortex being ripped open and requiring immediate repair. An important component of the cell’ s repair response is the formation of an actomyosin ring at the wound periphery to mediate its closure. Inhibition of linear actin nucleation factors and myosin result in a disrupted contractile apparatus and delayed wound closure. Here we show that branched actin nucleators function as a scaffold to assemble and maintain this contractile actomyosin cable. Removing branched actin leads to the formation of smaller circular actin-myosin structures at the cell cortex and slow wound closure. Removing linear and branched actin results in failed wound closure. Surprisingly, removal of branched actin and myosin results in the formation of parallel linear actin filaments that undergo a chiral swirling movement to close the wound. These results provide insight into actin organization in contractile actomyosin rings and uncover a new mechanism of wound closure.

**Summary:** Hui et al. find that branched actin is required during cell wound repair to serve as a scaffold to anchor the contractile actomyosin cable at the wound periphery. Inhibition of branched actin and myosin results in parallel linear filaments that swirl to close the wound, uncovering a new mechanism for cell wound repair.

## INTRODUCTION

All cells in the body undergo varying levels of physiological or environmental stresses—in response to normal daily events or as a result of diseases leading to fragile cells such as muscular dystrophy or diabetes—that can cause damage to the cell cortex (plasma membrane and underlying cortical cytoskeleton) (Cooper and McNeil, 2015; Galan and Bliska, 1996; Howard et al., 2011). This damage initiates a cell’ s rapid repair programs to minimize further damage, prevent infection or cell death, and restore homeostasis (Nakamura et al., 2018; Sonnemann and Bement, 2011; Velnar et al., 2009). Cell wound repair is highly conserved and is comprised of three main events: 1) rapid plasma membrane resealing; 2) dynamic cytoskeletal reorganization for wound closure; and 3) membrane and cytoskeleton remodeling after wound closure to restore the cell cortex to its unwounded state. The second of these major events involves the formation, then constriction, of a Rho family GTPases-regulated actin ring at the wound edge (Bement et al., 1999; Nakamura et al., 2018; Sonnemann and Bement, 2011; Yumura et al., 2014).

Rho family GTPases (Rho, Rac, Cdc42) are recruited to wounds in distinct spatiotemporal patterns (Abreu-Blanco et al., 2014; Golding et al., 2019). These proteins, through their downstream effector proteins, recruit actin to the wound where it forms a dense actin ring bordering the wound edge (Abreu-Blanco et al., 2011a; Abreu-Blanco et al., 2014; Nakamura et al., 2017). The subsequent constriction of this actin ring to pull the wound closed can be accomplished through different mechanisms. *Xenopus* oocytes can translocate the actin ring through myosin-independent actin treadmilling: the continuous assembly of actin at the inner edge and its disassembly at the outer edge of the actin ring (Burkel et al., 2012). In the *Drosophila* cell wound model, the actin ring contracts in a myosin-dependent manner to close the wound (Abreu-Blanco et al., 2011a; Abreu-Blanco et al., 2011b; Nakamura et al., 2018).

Downstream effectors of Rho family GTPases include both linear and branched actin nucleation factors. In general, linear actin nucleation factors function downstream of Rho, whereas branched actin nucleators function downstream of Rac and Cdc42 (Campellone and Welch, 2010; Goley and Welch, 2006). Linear nucleation factors include Diaphanous-related formins (DRFs) that have been shown to affect actin ring formation in cell wound repair (Abreu-Blanco et al., 2014). Branched actin nucleation factors include the Wiskott-Aldrich Syndrome (WAS) family proteins WASp, SCAR/WAVE, and WASH (cf. (Takenawa and Suetsugu, 2007)). WAS family proteins polymerize branched actin through interaction with the Arp2/3 complex, and often function to regulate dynamic actin organizations (Alekhina et al., 2017; Burianek and Soderling, 2013; Campellone and Welch, 2010; Massaad et al., 2013; Millard et al., 2004; Takenawa and Suetsugu, 2007).

Here we show that WAS family members exhibit distinct spatiotemporal recruitment patterns to the wound edge, as well as distinct wound closure phenotypes using the *Drosophila* cell wound repair model. Using super-resolution microscopy we find that different actin architectures are formed around the wound edge when we independently knockdown WASp, SCAR, or Wash. We find that WASp contributes to actin filament orientation as the actin filaments making up the actin ring exhibit a significant shift from random to anterior-posterior alignment. Additionally, in the removal of branched actin through knockdown of Arp3, excessively long linear actin filaments are observed suggesting that branched actin serves as a scaffold for linear actin filaments. Lastly, we uncovered a new mechanism of wound closure when the cell is unable to form a proper contractile apparatus wherein parallel linear actin filaments spiral inward to close the wound. Thus, our results highlight the complex interplay required between linear and branched actin nucleation factors to facilitate proper actin ring architecture formation and dynamic closure in cell wound repair.

## RESULTS

### Rac knockdown suggests a role for branched nucleation factors in cell wound repair

All three Rho family GTPases are required non-redundantly for cell wound repair. In particular, Rac is needed for the recruitment of actin to the wound edge, while Rho1 and Cdc42 are crucial for the formation and stability of the actin ring (Abreu-Blanco et al., 2014; Benink and Bement, 2005; Moe et al., 2021). Interestingly, some actin organization persisted at the wound edge in embryos where Rho1 or Diaphanous (Dia; *Drosophila* formin protein and Rho1 downstream effector) were knocked down (Abreu-Blanco et al., 2014), consistent with roles for Rac and/or Cdc42 in the repair process.

As *Drosophila* has three Rac genes (Rac1, Rac2, and Mtl), we modulated Rac activity by treating embryos with the pan Rac inhibitor NSC 23766 (Abreu-Blanco et al., 2014; Gao et al., 2004). Wounds were generated by laser ablation on the lateral side of nuclear cycle 4-6 *Drosophila* syncytial embryos expressing an actin reporter (sGMCA; (Kiehart et al., 2000)) (see Methods). In control embryos, actin accumulates in a highly enriched actomyosin ring bordering the wound edge and an elevated actin halo encircling the actin ring (Fig. 1A-C, G-I; Video 1). As shown previously, Rac inhibition results in an overexpansion of wounds, severely reduced recruitment of actin to the wound edge, and slower wound closure (Fig. 1D-I; Video 1) (Abreu-Blanco et al., 2014). These phenotypes suggest a role for branched actin nucleation factors as downstream effectors of Rac (and Cdc42) in this process.

**Figure 1.**
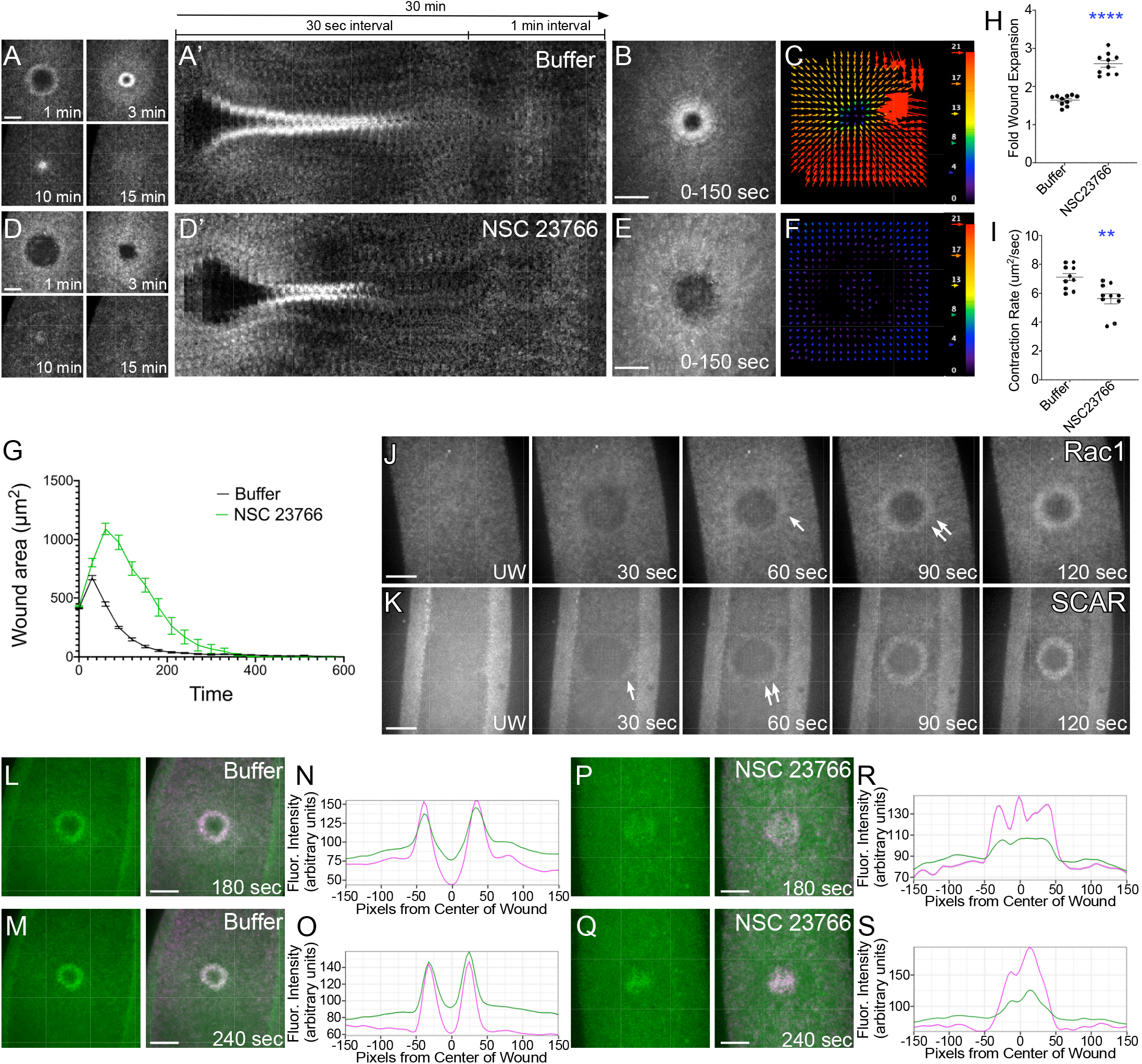
SCAR exhibits Rac independent recruitment patterns. **(A-F)** Confocal projection micrograph from time-lapse imaging of embryos expressing an actin marker (sGMCA) during cell wound repair in control (buffer only; A-B) or NSC 23766 injected (D-E). (A’, D’) Kymographs across the wound area in A and D, respectively. (B, E) XY max projection micrograph at 0-150s post wounding showing cortical flow of actin. (C, F) Vector maps from PIV (Particle Image Velocimetry) analysis depicting actin flow from 60-90s for A-B and C-D, respectively. **(G)** Quantification of the wound area over time for control (buffer injected) and NSC 23766 injected. **(H-I)** Quantification of wound fold expansion (H) and wound closure speed (I). **(J-K)** Recruitment of Rac1 and SCAR to cell wounds. Confocal XY projection micrographs of embryos coexpressing mCherry-Rac1 and sGMCA (J) or GFP-SCAR and sStMCA (K). **(L-M)** Confocal XY projection micrograph of embryos coexpressing GFP-SCAR, sStMCA, and injected with buffer at 180s (L) and 240s (M). **(N-O)** Fluoresence intensity (arbituary units) profiles across the wound area in L and M, respectively, at the times indicated. **(P-Q)** Confocal XY projection micrographs of embryos coexpressing GFP-SCAR, sStMCA, and injected with NSC 23766 at 180s (P) and 240s (Q). (**R-S**) Fluoresence intensity (arbituary units) profiles across the wound area in P and Q, respectively. Scale bars: 20μm. Error bars represent ± SEM. Unpaired student’ s t tests were performed in (H-I).

### SCAR is recruited to cell wounds in the absence of Rac

Consistent with its requirement for cell wound repair, Rac1 is recruited to the wound edge. As shown previously, wounding embryos expressing fluorescently tagged Rac1 (ChFP-Rac1) or Rac2 (GFP-Rac2) under the control of their endogenous promoters results in the strong accumulation of ChFP-Rac1 or GFP-Rac2 in a ring encircling the wound and in a less intense concentric ring corresponding to the actin ring and halo regions, respectively (Fig. 1J; Fig. S1A) (Abreu-Blanco et al., 2014; Nakamura et al., 2017). As Rho family GTPases have a one-to-one correspondence with WAS family proteins (Cdc42>WASp; Rac1>SCAR/WAVE; Rho1>Wash), we expected SCAR to show similar spatial and slightly delayed temporal recruitment to wounds as Rac1. We expressed GFP-tagged SCAR (GFP-SCAR) under the control of its endogenous promoter and find that it is recruited to wounds earlier than that observed for Rac1 and only in a ring encircling the wound (Fig. 1K). Thus surprisingly, Rac1 and SCAR recuitment to wounds is spatially and temporally distinct (Fig. 1J-K).

To futher investigate the relationship between Rac and SCAR in this context, we asked if recruitment of SCAR to cell wounds requires Rac. We examined GFP-SCAR recruitment to wounds in embryos injected with NSC 23766, a potent Rac inhibitor (Fig. 1L-S). Consistent with their different spatiotemporal recruitment patterns, GFP-SCAR was still recruited to wounds in these Rac inhibited embryos. Our results indicate that SCAR function is Rac-independent in the context of cell wound repair.

### WAS family members have distinct spatial and temporal recruitment patterns to cell wounds

To determine if and how other WAS family members are recruited to cell wounds, we expressed GFP-tagged WAS proteins, along with the sStMCA actin reporter (Fig. 2A; Video 2) (Nakamura et al., 2020). Upon laser injury, all WAS family members are recruited to wounds, albeit in distinct spatiotemporal recruitment patterns (Fig. 2B-G; Video 2). WASp is recruited to the wound with a peak accumulation at 90s, where it transiently colocalizes with the actin ring before dispersing by 240s post injury (Fig. 2B-B’, E; Video 2). SCAR recruitment overlaps with the actin ring (Fig. 2C-C’, F; Video 2). However, the peak recruitment of SCAR is much later (∼240 sec) than that of WASp (90 sec) or Wash (∼180 sec). Wash was more uniformly expressed throughout the embryo during the wound repair process and recruited less strongly to the wound edge compared to WASp and SCAR (Fig. 2D-D’, G; Video 2). Thus, the three WAS family members are recruited to the wound in different dynamic spatial and temporal patterns suggesting that they have distinct functions during cell wound repair.

**Figure 2.**
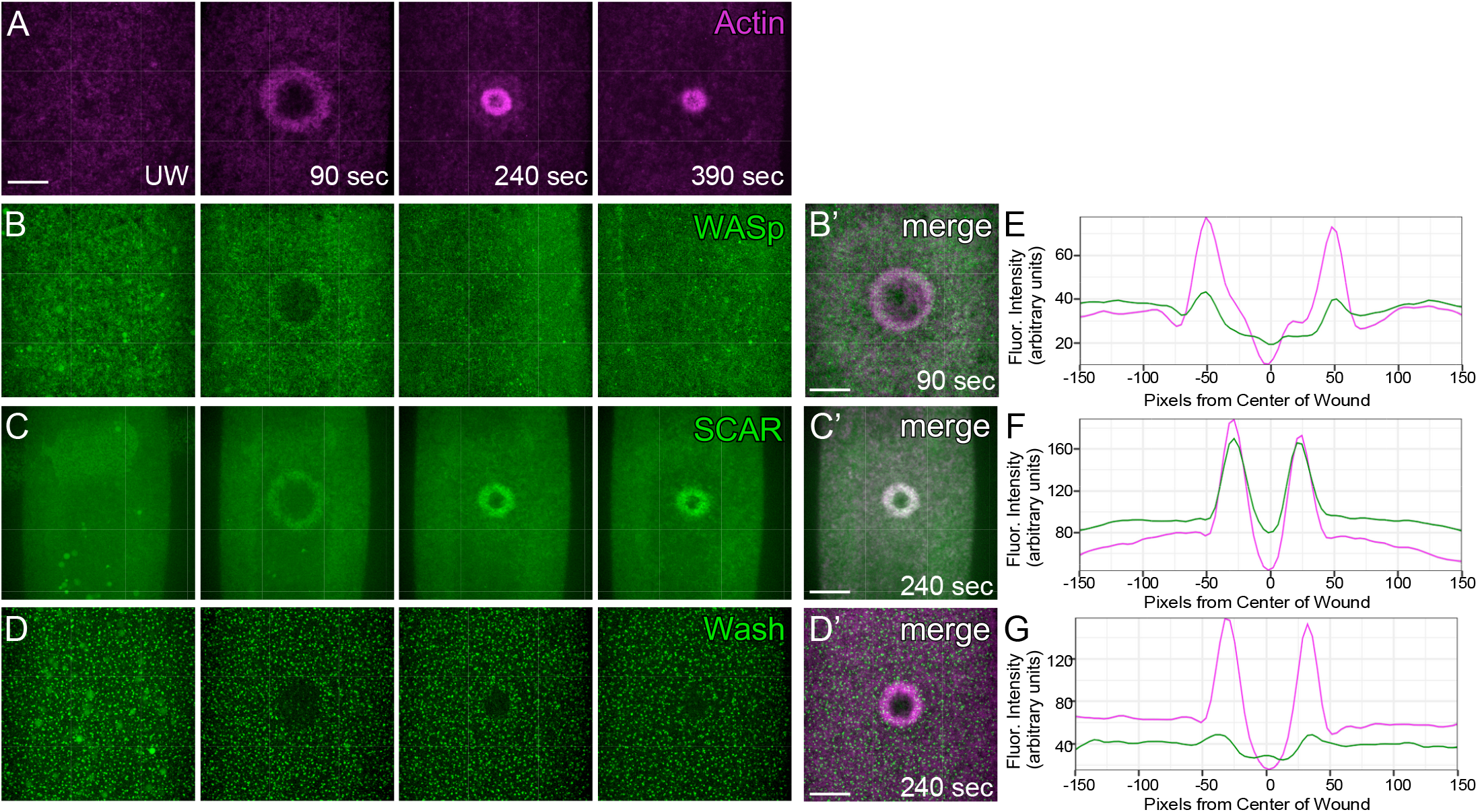
WASp, SCAR, and Wash exhibit distinct spatiotemporal recruitment patterns during cell wound repair. **(A-D’)** Confocal XY projection micrographs from time-lapse imaging of embryos co-expressing the sStMCA actin reporter (A-A’) with GFP-WASp (B-B’), GFP-SCAR (C-C’), or GFP-Wash (D-D’) at unwounded (UW), 90s, 240s, and 390s. **(E-G)** Fluoresence intensity (arbituary units) profiles across the wound area in B’ -D’, respectively. Scale bars: 20μm.

### Knockdown of WAS family members results in distinct defects in wound healing dynamics

Following the differences observed in the spatiotemporal recruitment patterns of the WAS family members, we expected each WAS subfamily to exhibit distinct effects on cell wound repair. To investigate this, we expressed the sGMCA actin reporter (Kiehart et al., 2000) in individual WAS family knockdown backgrounds (Fig. 3; Fig. S1B-E’ ; Video 3). Knockdown embryos were generated by expressing RNAi constructs in the female germline using the GAL4-UAS system (Brand and Perrimon, 1993; Rorth, 1998), using two independent RNAi lines for each WAS family member (Fig. S1F). Distinct cell wound repair phenotypes were observed in embryos lacking each of the WAS proteins. WASp knockdown resulted in minimal actin accumulation around the wound with a smaller actin ring width (WASp: 3.60±0.18µm, n=11; control: 5.62±0.24µm, n=13; p<0.0001), lower actin ring intensity (WASp: 1.20±0.02, n = 11; control: 2.271±0.13, n=13; p<0.0001), and wound overexpansion (WASp: 2.09±0.08 fold, n=11; control: 1.64±0.05 fold, n=13; p<0.001) (Fig. 3A-B’, F-G, K-N; Fig. S1B-B’ ; Video 3). Interestingly, knockdown of WASp did not show a significant difference in contraction rate (6.24±0.58 µm^2^/sec, n=11) compared to control (6.99±0.46 µm^2^/sec, n=13). Wounds generated in WASp knockdown fail to close completely within the 30min timelapse and exhibited loose actin accumulation at the center of the wound that is not observed in control embryos (Fig. 3A-B’, G, N; Fig. S1B-B’ ; Video 3).

**Figure 3.**
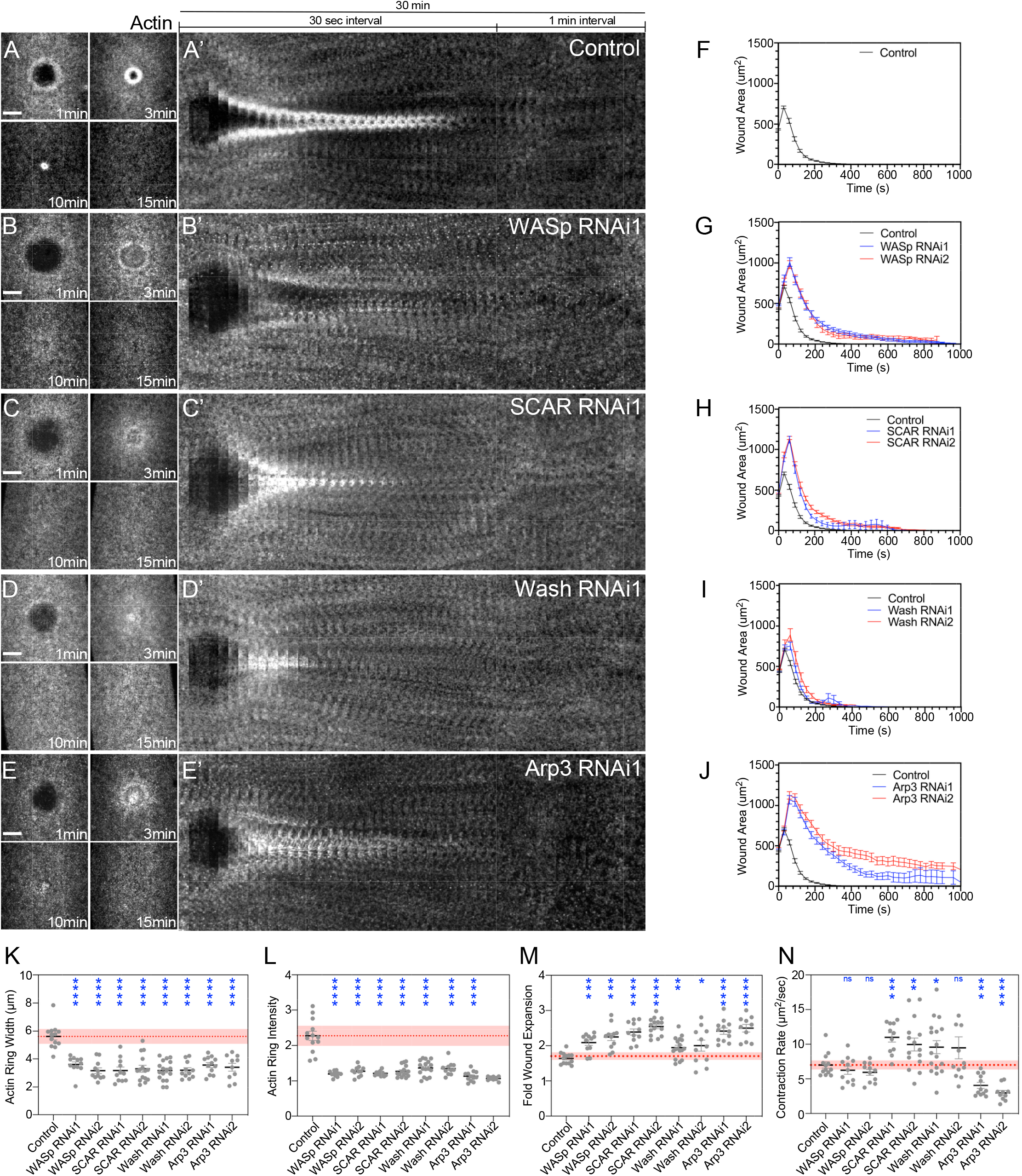
Branched actin is required for cellular wound repair and knockdown of each WAS family member exhibits non-redundant phenotypes. **(A-E’)** Actin dynamics (sGMCA) during cell wound repair: control (Vermilion RNAi; A), WASp RNAi (B), SCAR RNAi (C), Wash RNAi (D), and Arp3 RNAi (E). (A’ -E’) XY kymograph across the wound area depicted in A-E, respectively. **(F-J)** Quantification of wound area over time. **(K-N)** Quantification of actin ring width (K), average actin intensity (L), wound fold expansion (M), and wound closure rate (N). Scale bar: 20µm. Black line and error bars represent mean ± SEM. Red dotted line and square represent mean ± 95% CI from control. Unpaired Student’ s *t* tests were performed in K-N. * is p<0.05, ** is p<0.01, *** is p<0.001, **** is p<0.0001, and ns is not significant.

SCAR knockdown increased the wound contraction rate (10.99±0.67 µm^2^/sec, n=11; p<0.001) (Fig. 3C-C’, H, N; Fig. S1C-C’ ; Video 3). Similar to WASp, SCAR knockdown also resulted in significantly smaller actin ring widths (3.17±0.25µm, n= 11; p<0.0001), lower actin ring intensity (1.22±0.02, n=11; p<0.0001), and severe wound overexpansion (2.39±0.09 fold, n=11; p<0.0001) compared to control wounds (Fig. 3C-C’, H, K-N; Fig. S1C-C’ ; Video 3). Wash knockdown wounds exhibited a slight overexpansion phenotype (1.95±0.07, n=17; p<0.01) and a faster contraction rate (9.58±0.93 µm^2^/sec, n=17; p<0.05) (Fig. 3D-D’, I, K-N; Fig. S1D-D’ ; Video 3). Similar to WASp and SCAR, Wash knockdown also exhibited smaller actin ring width (3.17±0.17 µm, n=17; p<0.0001), reduced actin ring intensity (1.37±0.05, n=17; p<0.0001), and actin accumulation at the center of the wound. These results demonstrate that all three WAS family members are required for cell wound repair, and that they exhibit both overlapping and non-redundant roles during the repair process.

### Knockdown of Arp3 results in impaired branched actin nucleation

WAS proteins activate the Arp2/3 complex by facilitating the presentation of a G-actin monomer that works with Arp2 and Arp3 subunits to begin branched actin nucleation. Since the Arp2/3 complex is needed for actin nucleation activity of all WAS proteins, we expected that knockdown of Arp3 would mimick the combined loss of Wasp, SCAR, and Wash, thereby eliminating the majority of branched actin nucleation. As expected, Arp3 knockdown results in profound effects on cell wound repair: significantly less actin accumulation at the wound edge (1.14±0.04, n=12; p<0.0001), smaller actin ring width (3.56±0.18 µm, n=12; p<0.0001), and premature dissociation of actin ring structures from the surrounding actin cortex (Fig. 3E-E’ ; J, K-N; Fig. S1E-E’ ; Video 3). Additionally, Arp3 knockdown embryos also exhibit greater wound overexpansion (2.41±0.09 fold, n=12; p<0.0001) and a significantly slower contraction rate (4.06±0.44 µm^2^/sec, n=12; p<0.001) (Fig. 3M-N). Thus, Arp3 knockdown exhibits a combination of cell wound repair phenotypes observed in WASp, SCAR, and Wash knockdowns individually. Taken together, WAS proteins and their co-factor Arp2/3 play non-redundant and essential roles in cell wound repair, likely by influencing the spatiotemporal patterns of branched actin.

### WAS family coordinate actin ring filament architecture

Since WAS proteins and Arp2/3 influence the formation of branched actin and/or its assembly into different architectures, we performed time lapse imaging of actin dynamics following injury using super-resolution microscopy in WAS and Arp3 knockdowns. We captured the spatiotemporal differences in actin filament organization at the wound edge at both 50% and 70% wound closure (Fig. 4). Consistent with previous measurements of actin ring intensity (Fig. 3L), the actin ring in control embryos has a greater density of actin recruitment (Fig. 4A-A’“). Interestingly, knockdown of the WAS proteins resulted in clear polarization of actin filaments around the wound (Fig. 4B’ - D’”) unlike those observed in control wounds where they are more randomly oriented (Fig. 4A’ - A’”). Reduction of branched actin by Arp3 knockdown exhibited the most pronounced phenotype with the formation of circular actin structures in the unwounded state (Fig. 4E), limited actin recruitment around the wound, and long linear filaments at the center and around the wound (Fig. 4E’ -E’”). In contrast, knockdown of the major linear actin nucleator *Diaphanous* (Dia) did not exhibit the circular actin structures (Fig. 4F) or the polarized linear filaments (Fig. 4F’ -F’”) observed in WAS and Arp3 knockdowns.

**Figure 4.**
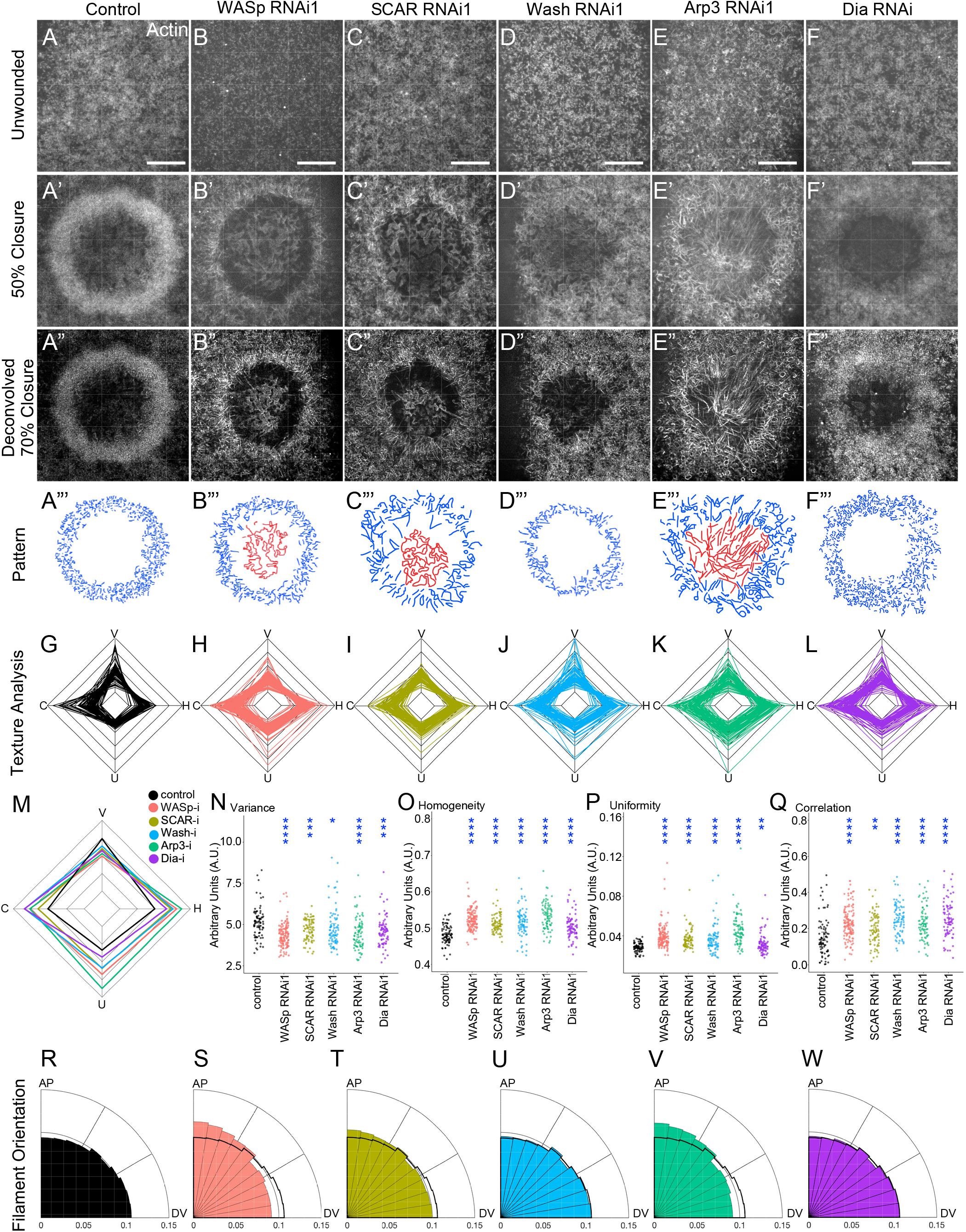
Individual knockdown of WAS family members, Arp3, and Dia exhibit different actin architectures. **(A-F’ ‘)** Super-resolution confocal micrographs from embryos expressing the sGMCA actin reporter: control (vermilion RNAi; A-A’ ‘), WASp RNAi (B-B’ ‘), SCAR RNAi (C-C’ ‘), Wash RNAi (D-D’ ‘), Arp3 RNAi (E-E’ ‘), and Dia RNAi (F-F’ ‘). (A-F) Unwounded actin cortex: control (vermilion RNAi) (A), WASp RNAi (B), SCAR RNAi (C), Wash RNAi (D), Arp3 RNAi (E), and Dia RNAi (F). (A’ -F’) Single z-plane micrographs of the injury site at 50% of max wound area from embryos shown in A-F, respectively. (A’ ‘ -F’ ‘) Single z-plane deconvolved images of the injury site at 70% of max wound area of embryos shown in A-F, respectively. **(A’ ‘ ‘ -F’ ‘ ‘)** Hand traced drawings showing actin filament organization in A’ ‘ -F’ ‘, respectively (blue: actin ring, red: actin internal to wound edge). **(G-L)** Radar charts depicting calculated Haralick features (Variance (V), Homogeneity (H), Uniformity (U), and Correlation (C)) of cropped actin ring regions from control (vermilion RNAi) (G), WASp RNAi (H), SCAR RNAi (I), Wash RNAi (J), Arp3 RNAi (K), and Dia RNAi (L). **(M)** Radar chart of average Haralick features for each genotype from G-L. **(N-Q)** Dot plots of Haralick features used in G-H. **(R-W)** Radial histograms showing the distribution of actin filament orientations ranging from Anterior-Posterior (AP) bias to Dorsal Ventral (DV) bias of control (vemilion RNAi) (R), WASp RNAi (S), SCAR RNAi (T), Wash RNAi (U), Arp3 RNAi (V), and Dia RNAi (W). Scale bar: 10µm. Error bars represent mean ± SEM. Unpaired Student’ s *t* tests were performed in N-Q. * is p<0.05, ** is p<0.01, *** is p<0.001, **** is p<0.0001.

To visualize the different actin organizations within the actomyosin ring, single z-slice images of wounds (0.25-0.5 µm below the cell cortex) at 70% closure were deconvolved and used for further analyses (see Methods; Fig. 4A”-F”). To quantify the bulk actin architectural differences, we performed texture analysis using Haralick features, including Variance/contrast: intensity between pairs of pixels; Correlation: linear dependency; Uniformity: proficient order of a whole image; and Homogeneity: tightness of pixel intensity distribution (Nakamura et al., 2020). Plotting these Haralick features on a radarchart, we observe a consistent pattern of actin ring architecture at wounds in WAS knockdown embryos (Fig. 4H-K, M) that differ significantly from the architecture observed in control wounds (Fig. 4G, M). Similarly, the Haralick features indicate an altered actin architecture at wounds in Dia knockdowns (Fig. 4L-M). All knockdown wounds exhibited significant decrease in variance compared to control (control: 5.14±0.11, n=80; WASp: 4.34±0.06, n=136, p<0.0001; SCAR: 4.69±0.07, n=80, p<0.001; Wash: 4.80±0.13, n=80, p<0.05; Arp3: 4.46±0.11, n=80, p<0.0001; Dia: 4.60±0.10, n=80, p<0.001) (Fig. 4N). Additionally, the actin organization of all knockdown wounds exhibited a significant (p<0.0001) increase in homogenity (control: 0.48±0.003; WASp: 0.52±0.002; SCAR: 0.51±0.003; Wash: 0.52±0.004; Arp3: 0.53±0.004; Dia: 0.50±0.004) and correlation (control: 0.16±0.01; WASp: 0.23±0.007; SCAR: 0.20±0.01, p<0.01; Wash: 0.25±0.01; Arp3: 0.23±0.01; Dia: 0.26±0.01) (Fig. 4O-P). When measuring the uniformity of the images, each WAS protein and Arp3 exhibited significant (p<0.0001) increase (control: 0.029±0.001; WASp: 0.04±0.001; SCAR: 0.04±0.001; Wash: 0.04±0.002; Arp3: 0.05±0.002), whereas Dia exhibited a marginal increase (0.032±0.001, p<0.01) (Fig. 4Q). These results indicate that knockdown of either branched or linear actin causes severe, albeit distinct, disruptions to the actin filament architecture of the contractile actomyosin ring at the wound edge.

### WASp is required for actin filament orientation at the wound edge

To further differentiate the roles of each WAS protein, we conducted actin filament orientation analyses by measuring the magnitude and direction of intensity gradients of actin filaments/bundles at the wound edge and expressing them as the ratio of filament orientations between 80°-90° (anterior-posterior alignment; AP) and 0°-10° (dorsal-ventral alignment; DV) (orientation bias; see Methods) (Fig. 4R-W). Control actin rings exhibited fairly uniform distribution of actin filaments (Fig. 4R) and an orientation bias of 0.89±0.02 (n=10). In contrast, WASp and Arp3 knockdown exhibited a shift towards AP orientation (WASp: 1.61±0.26, n=17, p<0.05; Arp3: 1.26±0.16, n=10, p<0.05) (Fig. 4S, V, X). Actin filaments in SCAR and Wash knockdown did not exhibit statistically significant (p>0.05) actin filament orientation shifts (SCAR: 1.038±0.08, n=10; Wash: 0.87±0.04, n=10) (Fig. 4T-U). Dia knockdowns do not exhibit an actin filament orientation bias (0.89±0.01, n=10) (Fig. 4W). Our results indicate critical roles for WASp and Arp3 in maintaining proper actin filament orientation at the wound edge. The lack of a clear orientation bias in SCAR and Wash knockdown highlights their distinct functions in cellular wound repair compared to WASp.

### Circular actin structures associate with myosin and Dia

Upon removal of branched actin (Arp3 knockdown), the unwounded cell cortex exhibits a striking accumulation of 0.6-1.3µm diameter circular actin filament structures (actin circles) that get incorporated into the actin ring upon wounding (Fig. 5A-C). Although some of these actin circles are present in the cortex of control embryos, embryos lacking branched actin have significantly more actin circle structures per region analyzed (control: 0.2±0.09, n=20; Arp3 RNAi: 11.8±1.1, n=20, p<0.0001) (Fig. 5A-C). To examine the potential role of these altered actin structures, we expressed either Myosin-StFP (Spaghetti Squash-mScarlet) or Dia-GFP, along with a fluorescent actin reporter (sGMCA or sStMCA, respectively), in an Arp3 RNAi background. We find that myosin associates with the majority of actin circles (67%; n=77) (Fig. 5D-H’), and does so in various patterns: accumulating in proximity to the actin circle (29%; n=33), filling the inside of the actin circle (13%; n=15), or only partially filling the actin circle (25%; n=29). In contrast, Dia was only observed to associate with 39% of the actin circles (Fig. 5I-K’). The association of myosin with the actin circles suggests an intriguing possibility wherein myosin bends and curls the untethered linear actin filaments into circular structures (Fig. 5L). Further, we find that both myosin and Dia associate with the long linear actin filaments that accumulate at the center of the wound (Fig. 5D-D’, I-I’), suggesting that the presence of Dia may prevent myosin from circularizing these actin filaments.

**Figure 5.**
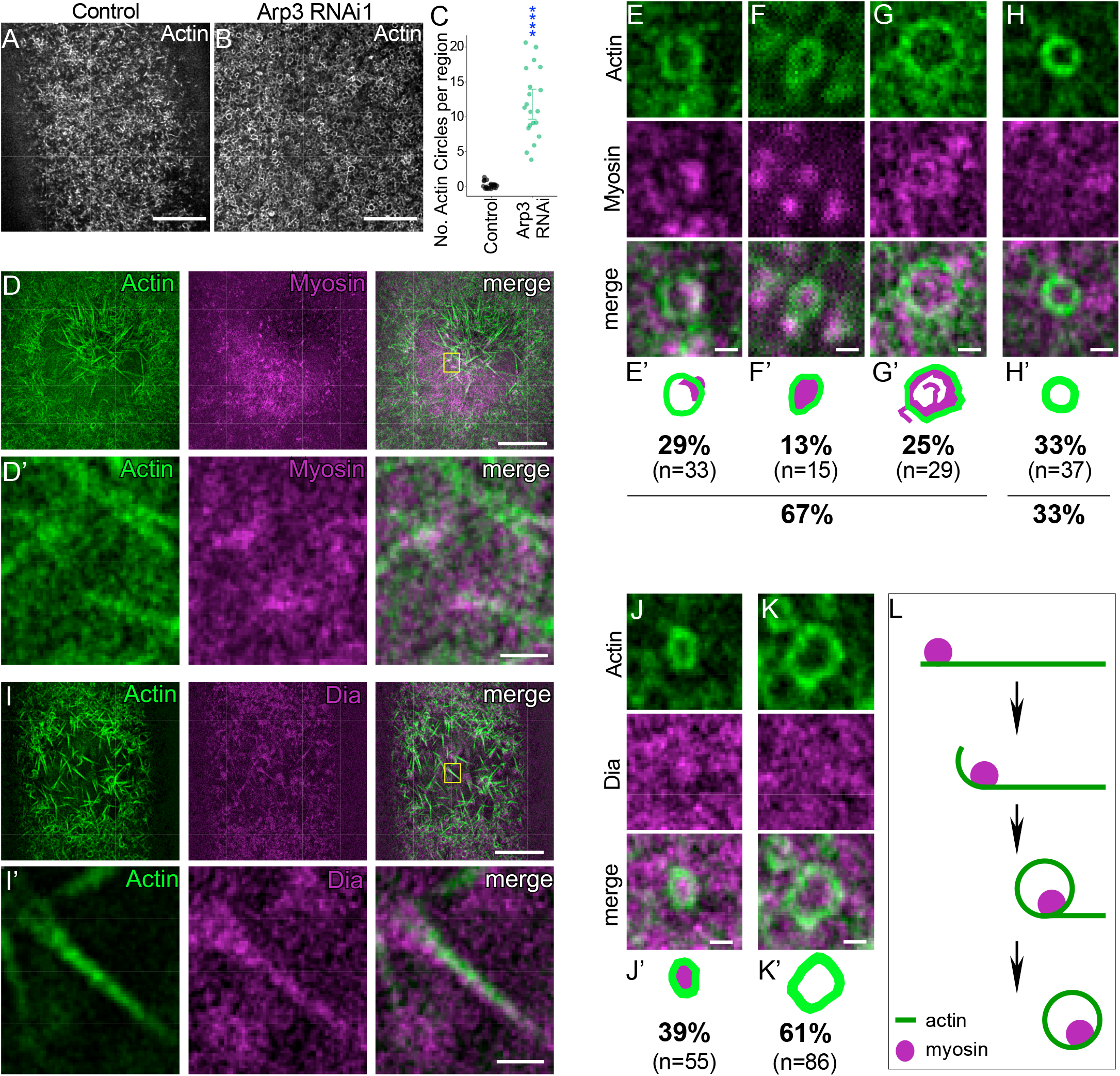
New circular actin structures form in the absence of branched actin. **(A-B)** Deconvolved single slice confocal micrographs of the unwounded cortex of embryos expressing the sGMCA actin reporter: control (vermilion RNAi) (A) and Arp3 RNAi (B). Scale bars: 10µm. (**C**) Quantification of the number of actin circles per region (380µm^2^) in A-B. (**D-D’**) Deconvolved single slice confocal micrographs of Arp3 RNAi embryos co-expressing the sGMCA actin and Myosin (spaghetti squash)-mScarlet reporters at 5min post wounding. Scale bar: 10µm. Higher magnification view of yellow highlighed region (D’). Scale bar: 1µm. (**E-H’**) Higher magnification view and quantifcation of actin circle and myosin associations observed. Drawings depicting actin circle and myosin association patterns shown in E-H, respectively (E’ -H’). Scale bars: 0.5µm. (**I-I’**) Deconvolved single slice confocal microgaphs of Arp3 RNAi embryos co-expressing the sStMCA actin and Dia-GFP reporters. Scale bar: 10µm. Higher magnification view of yellow highlighted region (I’). Scale bar: 1µm. (**J-K’**) Higher magnification view and quantification of actin circle and Dia association. Drawings depicting actin circle and Dia association (J”-K’). Scale bar: 0.5µm. (**L**) Model of actin circle formation in the absence of branched actin and association with myosin.

### Coordinated action of myosin with branched and linear actin is required for proper actomyosin ring assembly

Since both linear and branched actin nucleation factors are needed for cell wound repair, yet some actin organization still exists at the wound edge when either one is removed, we next determined what actin organization, if any, remained when linear and branched actin nucleation were removed simultaneously. To this end, we removed linear actin using either a Dia knockdown (major formin present during cell wound repair) or treatment with C3 exoenzyme (Rho inhibitior; affects several linear nucleators including formins and Spire) in an Arp3 knockdown (no branched actin) background (Fig. 6A-C’’). Unsurprisingly, removing linear actin by inhibiting Rho1 activity (C3 exoenzyme) in this context severely disrupted the actin cortex and resulted in very few discernable actin structures and failure to close wounds compared to Arp3 RNAi embryos injected with buffer (Fig. 6A-B’’, F), suggesting that branched and linear actin account for the majority, if not all, of the actin organization required at wounds. Inhibiting linear actin with Dia knockdown led to a significant decrease (p<0.0001) in the number of actin circles compared to Arp3 RNAi injected with buffer (buffer: 11.85±0.79, n=20; Dia RNAi: 5.92±0.67, n=24; C3: 0.75±0.25,n=20) (Fig. 6C-C’’, F), suggesting that Dia is required for actin circle formation.

**Figure 6.**
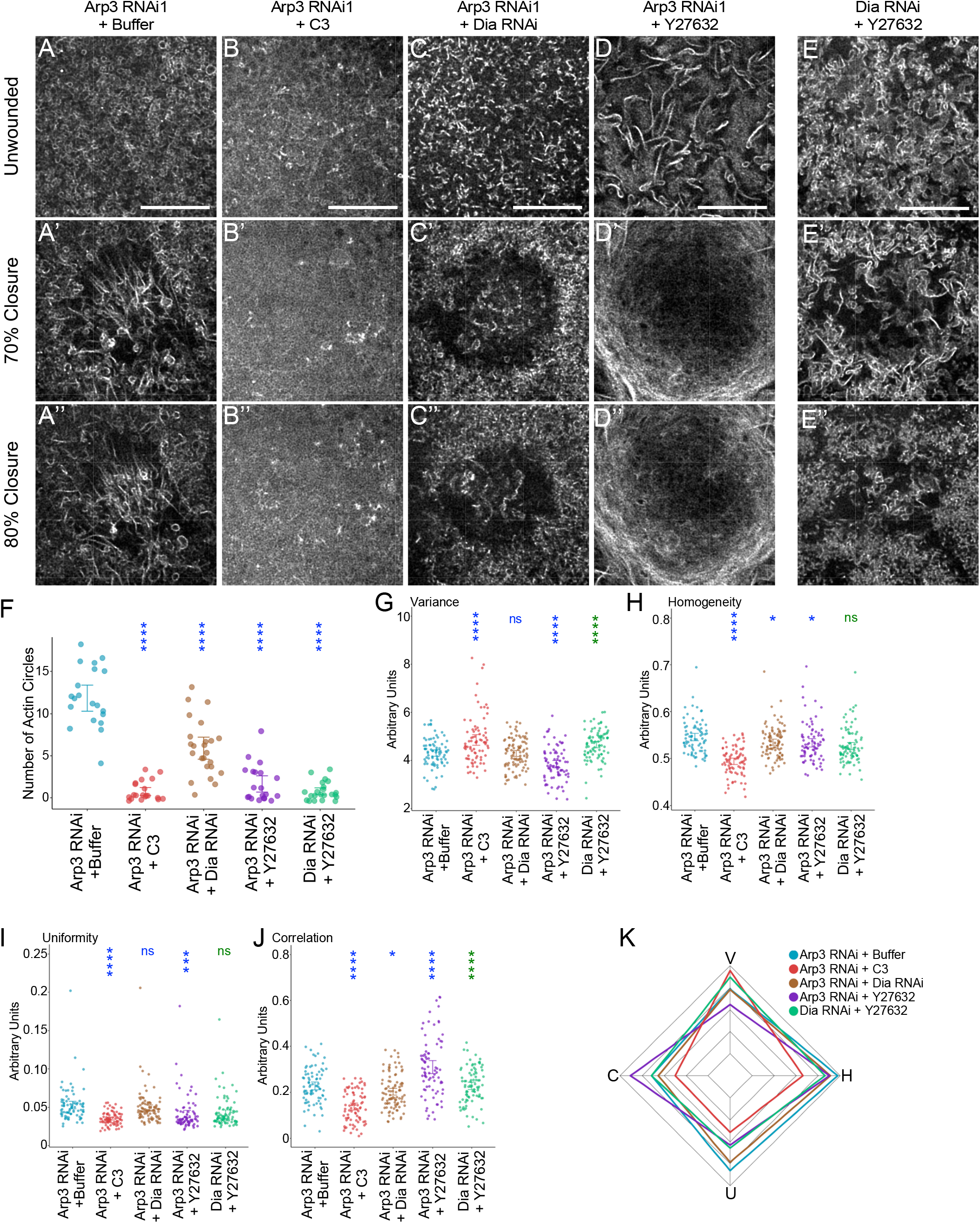
Both branched and linear actin are vital to wound repair. **(A-D’ ‘)** Super-resolution confocal micrographs of embryos co-expressing the sGMCA actin reporter and Arp3 RNAi that have been treated with Buffer only (control, A-A’ ‘), C3 exoenzyme (B-B’ ‘), Dia RNAi (C-C’ ‘), or Y27632 (D-D’ ‘). **(E-E’ ‘)** Super-resolution confocal micrographs of embryos co-expressing the sGMCA reporter, Dia RNAi, and treated with Y27632. (A’ -E’) Deconvolved single z-plane micrographs of the injury site at 70% of max wound area of embryos shown in A-D, respectively. (A’ ‘ -E’ ‘) Deconvolved single z-plane micrographs of the injury site at 80% of max wound area of embryos shown in A-F, respectively. **(F)** Quantification of the number of actin circles in a 300×300 pixel area of the unwounded cortex. **(G-J)** Dot plots of calculated Haralick features of cropped actin ring/wound edge regions. **(K)** Radar chart of average Haralick features for each genotype in G-J. Scale bar: 10µm. Error bars represent mean ± SEM. Unpaired Student’ s *t* tests were performed in F-J. * is p<0.05, ** is p<0.01, *** is p<0.001, **** is p<0.0001.

We next disrupted myosin activity in Arp3 knockdowns by treatment with the ROK inhibitor Y27632. Strikingly, this resulted in the presence of bundled linear actin filaments in the unwounded cortex (Fig. 6D-D’’). Inhibiting myosin activity also led to a significant reduction (p<0.0001) in the number of actin circles (buffer: 11.85±0.79, n=20; Y27632: 1.65±0.49, n=20) (Fig. 6D-D”, F), suggesting that myosin is also required for actin circle formation. To examine the role of linear actin in this large bundled actin phenotype, we disrupted myosin in a Dia knockdown background. The inhibition of linear actin and myosin led to an unwounded actin cortex consisting primarily of unbundled actin patches with scattered bundled linear actin filaments (Fig. 6E-E’’). This is consistent with branched actin serving as a scaffold for linear actin organization, whereas myosin is required for branched actin disassembly to prevent its aggregation. In addition, without linear actin or myosin, the number of actin circles are significantly reduced compared to buffer injected Arp3 RNAi (0.77±0.22, n=20, p<0.0001) (Fig. 6F).

To quantify the actin architectures in these contexts, we performed texture analysis of the wound edge at 70% closure (Fig. 6A’ -E’). Embryos treated with Y27632 or C3 exhibited a significant increase in variance compared to embryos treated with buffer (buffer: 4.31±0.07, n= 80; Y27632: 3.85±0.08, n= 80, p<0.0001; C3: 4.85±0.11, n=80, p<0.0001) (Fig. 6G). In contrast, Dia knockdown did not show a statistically significant change (4.29±0.06, n=96). Both Y27632 and C3 treatment exhibited a decrease in homogeneity (buffer: 0.55±0.004; Y27632: 0.53±0.005, p<0.05; C3: 0.49±0.003, p<0.0001) and uniformity (buffer: 0.05±0.003; Y27632: 0.04±0.003, p<0.001; C3: 0.03±0.001, p<0.0001) (Fig. 6H-I). Additional Dia knockdown exhibited a significant decrease in homogeneity (0.54±0.003, p<0.05), but not in uniformity (0.05±0.002, p>0.05). Lastly, both Dia knockdown and C3 treatment exhibited significant decrease in correlation (buffer: 0.23±0.01; Dia: 0.20±0.01, p<0.05; C3: 0.14±0.01, p<0.0001) (Fig. 6J). In contrast, embryos treated with Y27632 exhibited a significant increase in correlation (0.31±0.01, p<0.0001). Wound edges lacking linear actin and myosin exhibited a significant increase in variance (4.66±0.07, n=88, p<0.0001) and a significant decrease in correlation (0.23±0.01,n=88, p<0.0001) compared to wounds lacking branched actin and myosin. In contrast, uniformity (0.04±0.002, n=88) and homogeneity (0.53±0.004, n=88) of the actin architecture did not show statistical significance (p>0.05). Taken together, the differences observed among these Haralick features indicate the presence of different actin architectures/assemblies in these different knockdown contexts (Fig. 6K).

### New mechanism of wound closure in the absence of branched actin and myosin

Without branched actin to serve as a scaffold for building/tethering the actomyosin contractile apparatus, Arp3 knockdowns treated with Y27632 form bundled linear actin filaments in the unwounded cortex (Fig. 7A; Video 4). Upon wounding, these bundled linear actin filaments organize and align into striking parallel arrays around the wound edge, which then move in a coordinated fashion to reduce the wound area (actin swirling: 91%, n=11) (Fig. 7A-B). This movement of actin filaments is not observed in either Arp3 knockdown with buffer injection or in Dia knockdown with Y27632 injection (Fig. 7C). Parallel microtubule arrays have been shown to slide past one another bidirectionally to create cytoplasmic movements in various developmental contexts (Jolly and Gelfand, 2010; Lu et al., 2013; Lu et al., 2016). To identify whether the parallel actin filaments at the wound edge moved together or were sliding past one another in opposite directions, we expressed a photoconvertible actin marker (sM3MCA) to track actin filament movement. Intriguingly, the parallel linear actin filaments consistently move together in an anti-clockwise direction (100%, n=10) (Fig. 7D-E), suggesting specific chirality to this newly uncovered wound closure mechanism.

**Figure 7.**
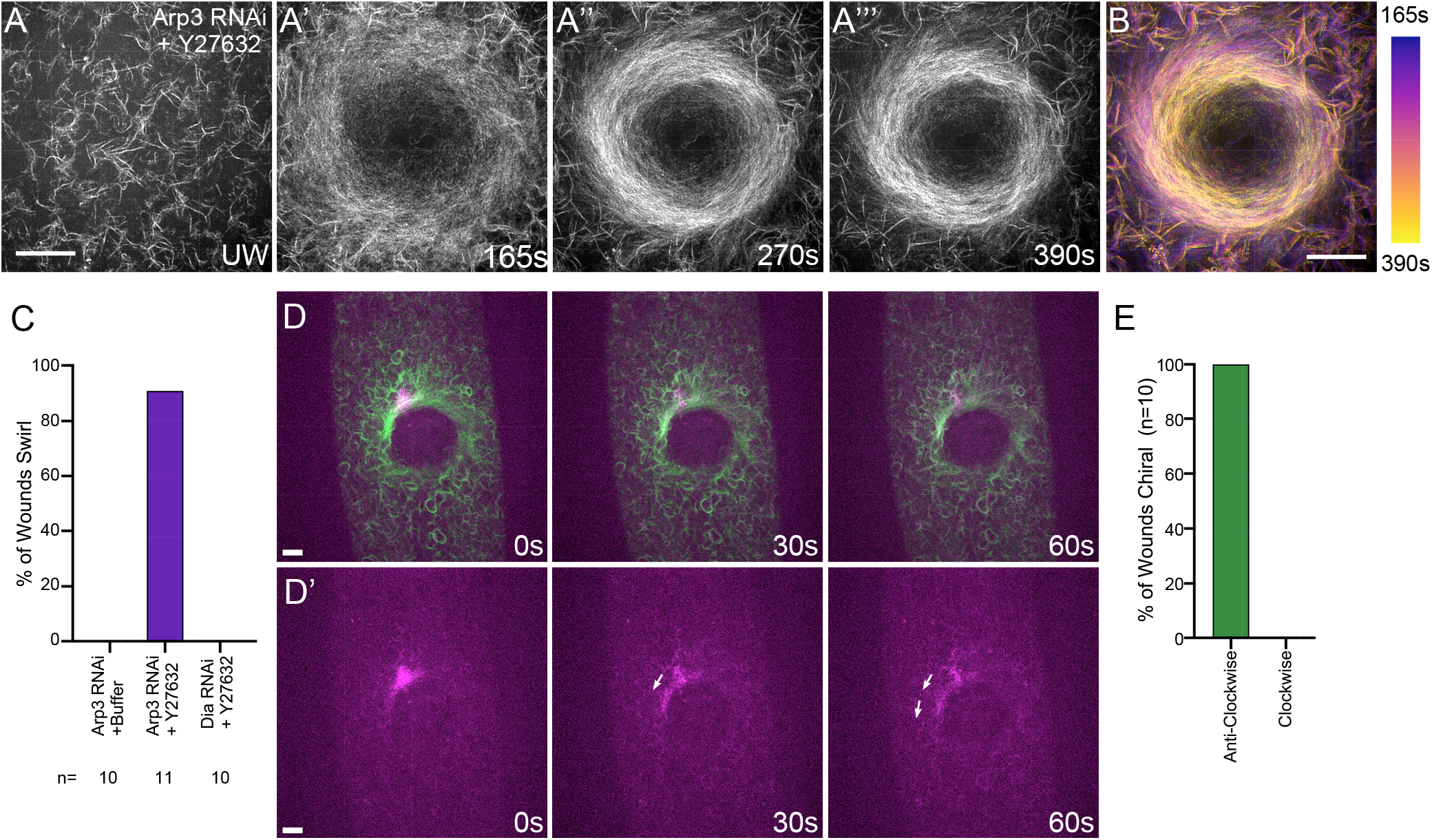
In the absence of branched actin and myosin, linear actin filaments swirl to close the wound. **(A-A’ ‘ ‘)** Confocal XY max projection micrographs from time-lapse imaging of an Arp3 RNAi embryo treated with Y27632 and expressing the sGMCA actin reporter at unwounded stage (A), 165s (A’), 270s (A’ ‘), and 390s (A’ ‘ ‘). **(B)** Temporal-colour coded projection of actin swirling at the wound depicted in A-A’ ‘ ‘ from 165s (purple) to 390s (yellow). **(C)** Quantification of percentage of wounds that exhibited actin filament swirling. **(D-D’)** Confocal XY max projection micrographs from time-lapse imaging of an Arp3 RNAi embryo expressing a photoconvertible actin reporter (sM3MCA). Photoconverted region showing counter-clockwise movement of actin filaments (arrows) is highlighted in D’. **(E)** Quantification of number of wounds that exhibited actin filament swirling in the anti-clockwise direction. Scale bar: 10µm.

## DISCUSSION

Cell wound repair is a highly conserved process that relies heavily on the formation, then dynamic translocation, of a contractile actomyosin ring at the wound periphery. To accomplish this, specific organizations of actin filaments (mixed orientation), non-muscle myosin, scaffolding proteins, and associated crosslinking proteins must be precisely assembled then undergo intricately coordinated changes as the ring pulls the wound closed. Here we show that successful cell wound repair requires both linear and branched actin networks for the proper assembly and function of the actomyosin ring. Some actin organization remains at the wound edge when either type of actin is knocked down leading to delayed wound closure, whereas all actin organization is lost when both are knocked down leading to failed wound repair. We find that the WAS family branched actin nucleation promoting factors (WASp, SCAR, and Wash) exhibit different spatiotemporal recruitment patterns to wounds and have both overlapping and non-redundant roles in the repair process. Interestingly, we find that WASp contributes to actin filament orientation within the actomyosin ring and the loss of branched actin via Arp3 knockdown results in the formation of numerous small actin circles in the cell cortex. Intriguingly, we find that the simultaneous lack of branched actin and myosin activity results in an unexpected actin organization wherein an array of parallel actin filament bundles undergoes concerted movement resulting in a chiral swirling inward to close the wound, uncovering a new mechanism for cell wound closure.

### Branched and linear actin coordinate proper actin architecture for cell wound repair

Many diverse cell processes such as cell division, motility, and shape changes require that actin filaments be organized into assemblies of optimized architecture, dynamics, and mechanical attributes to carry out their specific functions properly (Boiero Sanders et al., 2020; Campellone and Welch, 2010; Cheffings et al., 2016; Gautreau et al., 2021; Schwayer et al., 2016). We find that during cell wound repair the proper assembly and function of the actomyosin ring requires both branched and linear actin networks (Fig. 8A). This is consistent with observations that *in vitro* actomyosin rings consisting of only linear or only branched actin exhibit impaired contraction (Ennomani et al., 2016). Overlapping bundles of linear actin filaments in mixed orientation, along with non-muscle myosin motors and crosslinking proteins, make up the contractile actin cable that surrounds the wound periphery (Blanchoin et al., 2014; Cheffings et al., 2016; Schwayer et al., 2016). We find that branched actin networks contribute in several ways during cell wound repair: as a scaffold, to influence filament orientation and length, as a foundation to assemble and tether the actomyosin ring, and to aid in anchoring the cortical cytoskeleton to overlying plasma membrane. When branched actin is removed, linear actin bundles are unable to assemble into a functional cable, leading to the formation of small circles of actin and myosin. Conversely, we find that a cytoskeleton consisting primarily of branched actin without myosin results in the formation of dispersed amorphous actin structures (Fig. 8B). These structures are likely small aggregates of branched networks and are reminiscent of isotropic branched actin networks that are assembled in response to mutant Arp2/3 complexes, but are unproductive since they cannot organize into force-producing actin networks (Narvaez-Ortiz and Nolen, 2022).

**Figure 8.**
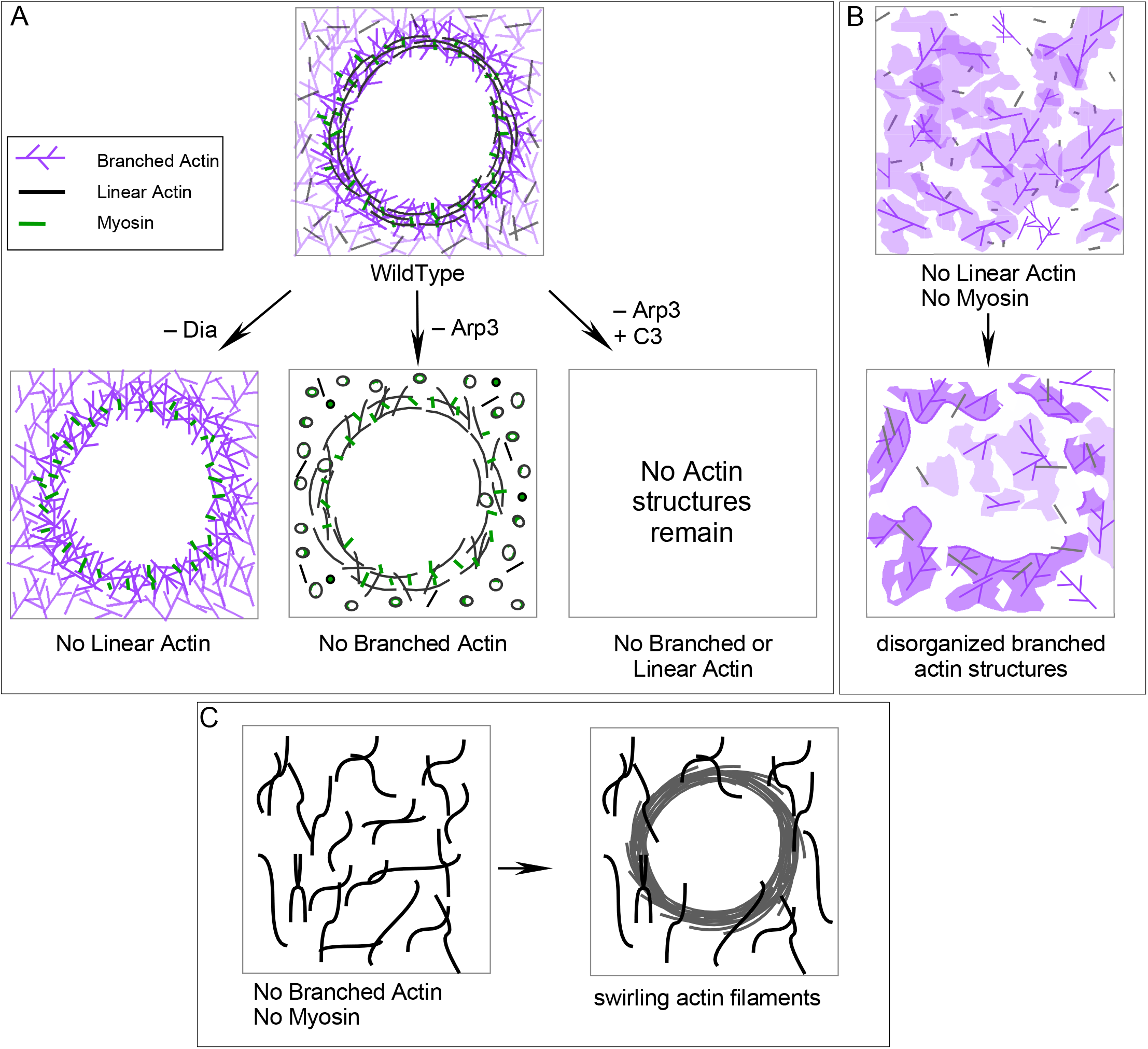
Model of branched and linear actin filament coordination during cell wound repair. **(A)** Branched actin acts as a structural scaffold to hold the linear actin cable at the wound edge. Without branched actin, subunits of the actin cable release from the wound edge and form actin circles due to myosin contraction and bending. Branched and linear actin are vitral for cell wound repair as no actin structures are formed upon Arp3 knockdowns treated with C3 injection. **(B)** The actin cortex forms large amorphous actin patches in the absence of linear actin and myosin (Dia RNAi + Y27632 treatment). **(C)** The cell forms large linear actin filaments and bundles in the absence of branched actin and myosin (Arp3 RNAi + Y27632 treatment). Intriguingly, upon wounding, these large linear actin filaments/bundles are organized at the wound edge and undergo a chiral swirling movement to close the wound.

While all WAS family members, along with Arp2/3, promote the generation of branched actin networks, specific family members can be called upon to perform specialized functions within the cell (Burianek and Soderling, 2013; Massaad et al., 2013). These non-redundant functions are often associated with the variable N-terminal domains of WAS family proteins that mediate different protein interactions leading to variations in the architecture of their actin assemblies (Gautreau et al., 2021; Molinie and Gautreau, 2018). Interestingly, WASp, Wash, and SCAR are all required for cell wound repair, where they exhibit different spatiotemporal recruitments and phenotypes.

Knockdown of WASp leads to a significant shift in actin filament orientation, highlighting the role of branched actin in bending and/or aligning linear actin filaments around the wound edge. Actin filament orientation is important for function, as myosin sliding along linear actin bundles for effective contraction requires a ring geometry with mixed actin filament polarities (Blanchoin et al., 2014; Cheffings et al., 2016). Our findings are also consistent with studies demonstrating a role for WASP in forming actin structures orthogonal to the cell’ s plasma membrane (Gaertner et al., 2022), as well as polarity during hemocyte (Stramer et al., 2005) and neutrophil (Brunetti et al., 2022) migration. Loss of branched actin also led to the formation of long linear actin filament bundles in the center of the wound. This is consistent with recent findings that the Arp2/3 complex is required to prevent excessive formin mediated linear actin nucleation (Chan et al., 2019).

SCAR knockdown exhibited a wound overexpansion phenotype, suggesting a potential role for branched actin in anchoring the plasma membrane to the underlying cortical cytoskeleton. We showed previously that E-cadherin functions in cell wound repair to anchor the actomyosin ring to the overlying plasma membrane in cell wound repair (Abreu-Blanco et al., 2011a). Interestingly, recent experiments in *C. elegans* have linked the WAVE/SCAR complex and associated branched actin to E-cadherin localization (Cordova-Burgos et al., 2021; Sasidharan et al., 2018). Alternatively, a recent study identified the orthogonal roles of WAVE1 and WAVE2 in regulating lamellipodial actin extension (Tang et al., 2020). Therefore, it is possible that *Drosophila* SCAR can additionally play a role in regulating the overall length of actin filaments in the actin ring. Thus, a complex and dynamic interplay of linear and branched actin is necessary for robust cell wound repair.

### Actin circle formation in the absence of branched actin scaffolding

In addition to affecting the actin architecture within the actin ring during wound repair, knockdown of Arp3 resulted in the formation of new circular actin structures throughout the cortex. These actin circles are associated with myosin. As the actomyosin ring does not form in the absence of anchoring to a branched actin scaffold, one possibility is that these actin/myosin circles are subunits of the actomyosin ring that are unable to assemble into the higher order structure. In this case, linear actin filaments may bend as a result of their association with actin binding proteins (Fassler et al., 2020; Harris et al., 2020; McCullough et al., 2008; Palani et al., 2021). Alternatively, a previous study demonstrated preferential branched actin nucleation on the convex side of bent actin filaments as a likely response to applied forces (Risca et al., 2012). Thus, it is possible that branched actin nucleation on the convex side of a filament can act to counter the bending forces to straighten the actin filament. It is also possible that without branched actin, linear filaments bend or encircle themselves due to myosin contraction. Consistent with this, myosin colocalizes with the actin circles and its inhibition abolishes actin circles and resulted in thicker linear actin bundles. We find that these actin circles are not observed in the cortex of WT embryos, suggesting that branched actin is presumably inhibiting the formation of this type of architecture. Thus, in the absence of branched actin, the delicate balance of actin binding proteins and branched and linear actin filaments results in the observed actin circles. Further investigations will be needed to elucidate molecular mechanisms behind the formation of actin circles and their functions in the cytoskeleton.

### A cell cortex comprised of linear actin filaments only exhibits a new mechanism of wound repair

The molecular mechanisms underpinning actin ring translocation during wound closure can be different depending on the context. *Xenopus* oocytes, for example, can use actin treadmilling to continuously assemble actin at the inner edge of the actin ring and disassemble actin at the outer edge to reduce the wound area (Burkel et al., 2012). *Drosophila* embryos require myosin to form a contractile actomyosin ring assembly to pull the wound closed (Abreu-Blanco et al., 2014). Surprisingly, we observe an unexpected wound repair mechanism when only linear actin filaments are available. When myosin activity and branched actin are removed, the remaining linear actin filaments form a parallel array that undergoes a chiral swirling movement wherein actin filaments move directionally to spiral inward and close the wound (Fig. 8C). This counter-clockwise actin filament swirling in cell wound repair is superficially similar to chiral cytoskeleton assemblies observed in other systems. Active chiral torques are generated to break left/right symmetry in the actomyosin cortex of *C. elegans* embryos and during actin cytoskeleton self-organization in cells with isotropic circular shape (Naganathan et al., 2014; Tee et al., 2015), as well as through the organization of formin-dependent linear actin filaments at immunological synapses (Murugesan et al., 2016). However, in all of these cases, the formation of the chiral cytoskeleton torques depend on myosin activity, whereas the chiral actin filament swirling during cell wound repair occurs in its absence. One class of proteins that been shown to enhance the contractility of myosin-independent (as well as myosin-dependent) actin rings *in vitro* are actin crosslinkers (Ennomani et al., 2016; Kucera et al., 2021). Loss of myosin and branched actin leads to a sparse actin network that consists of mainly larger bundles of actin, thus making actin crosslinkers a strong candidate to drive this phenotype. Future investigations will be required to elucidate the precise molecular mechanism leading to this chiral swirling phenotype.

Thus, while cells prefer to use a highly orchestrated arrangement of branched actin, linear actin, and myosin to efficiently close wounds, it can nevertheless co-opt whatever actin structures are available—branched actin only, linear actin plus myosin, or linear actin only—to close wounds (albeit more slowly). Future investigations will continue to unravel the complexities of these dynamic actin structures and identify the possible molecular triggers that dictate actin ring contraction, treadmilling, or filament swirling, as well as the roles of accessory actin binding proteins (i.e., crosslinkers) that can influence their spatiotemporal localizations, thus affecting the actin architecture and the overall mechanics of the cytoskeleton.

## Abbreviations List

Ch: mCherry fluorescent protein
Dia: Diaphanous
GFP: Green fluorescent protein
GLCM: Gray Level Cooccurrence Matrix
NC: Nuclear cycle
AP: Anterior-Posterior
DV: Dorsal-Ventral
sChMCA: spaghetti squash driven, mCherry fluorescent protein, moesin-α-helical-coiled and actin binding site
sfGFP: super folder green fluorescent protein
sGMCA: spaghetti squash driven, GFP, moesin-α-helical-coiled and actin binding site
sStMCA: spaghetti squash driven, mScarlet fluorescent protein, moesin-α-helical-coiled and actin binding site
sM3MCA: spaghetti squash driven, Maple3 fluorescent protein, moesin-α-helical-coiled and actin binding site
sqh: spaghetti squash
WAS: Wiskott-Aldrich Syndrome

## MATERIALS AND METHODS

### Fly stocks and genetics

Flies were cultured and crossed at 25°C on yeast-cornmeal-molasses-malt extract medium. Flies used in this study are listed in Table S1. RNAi lines were driven using the maternailly expressed GAL4-UAS driver, Pmatalpha-GAL-VP16V37.

An actin reporter, sGMCA (spaghetti squash driven, moesin-alpha-helical-coiled and actin binding site fused to GFP) reporter (Kiehart et al., 2000) or the mScarlet-i fluorescent equivalent, sStMCA (Nakamura et al., 2020), was used to follow wound repair dynamics of the cortical cytoskeleton.

In this study, we used an actin reporter + maternal GAL4 driver + *vermilion* RNAi (unrelated fly RNAi) as the control.

Localization patterns and mutant analyses were performed at least twice with independent genetic crosses and ≥10 embryos were examined. Images representing the average phenotype were selected for figures.

### GFP-tagged Scar reporter

To generate GFP-SCAR under its endogenous promoter, we first amplified full length Scar including the 5’ -UTR and the 3’ -UTR and cloned it into pBluescript using 5’ KpnI and 3’ SalI restriction sites. We then fused the 5’ -UTR, GFP, and Scar up to the internal XbaI site. This fragment was then fused from the XbaI internal site to the end of the 3’ -UTR and cloned into pCasper using 5’ KpnI and 3’ NotI restriction sites.

### Scarlet-tagged spaghetti squash knock-in (sqh-StFP)

sqh-StFP was generated using CRISPR-mediated homologous recombination. gRNA (5’ - TTACTGCTCATCCTTGTCCT-3’) was cloned into pCFD5 vector (Port et al., 2014) (Addgene #73914) and then the resulting vector was injected into attP2 site (BDSC #25710). Subsequently, third chromosome insertion transgenic pCFD5-sqh flies were crossed into w^-^ backgrounds. To generate the donor vector, mScarlet-i and 2kb homology arms were amplified then all three fragments were fused and cloned into the pBluescriptII KS(+) vector such that mScarlet-i was inserted just before the stop codon of the *sqh* gene. The donor vector was injected into embryos resulting from crossing nos-Cas9 (BDSC #78782 that had been crossed into a *w*^*-*^ background) and pCFD5-sqh flies. The mScarlet-i insertion was confirmed by sequencing and the sqh-StFP knock-in flies are viable and fertile.

### Photoconvertible actin reporter (sM3MCA)

To generate sM3MCA (spaghetti squash driven, mScarlet fluorescent protein, moesin-α-helical-coiled and actin binding site), we replaced mCherry sequence in sChMCA with mMaple3 (Wang et al., 2014) using standard PCR and cloning techniques.

### Embryo handling and preparation

Nuclear cycle 4-6 embryos were collected for 30min at 25°C and harvested at room temperature (22°C). Collected embryos were dechroionated by hand, mounted onto No. 1.5 coverslips coated with glue, and covered with Series 700 halocarbon oil (Halocarbon Products Corp.) as previously described (Abreu-Blanco et al., 2011a).

### Drug Injections

Pharmacological inhibitors were injected into NC4-6 staged *Drosophila* embryos, incubated at room temperature (22°C) for 5 min, and then subjected to laser wounding. The following inhibitors were used: C3 exoenzyme (1 mg/ml; Cytoskeleton, Inc), NSC23766 (50 mM; Tocris Bioscience), and Y27632 (60 mM; Tocris Bioscience). The inhibitors were prepared in injection buffer (5 mM KCl, 0.1 mM NaP pH6.8). Injection buffer alone was used as the control.

### Laser Wounding

All wounds were generated using a pulsed nitrogen N2 micropoint laster (Andor Technology Ltd.) set to 435nm and focused on the lateral surface of the embyro. A 16×16 µm circular region was set as the target site along the lateral midsection of the embryo, and ablation was controlled by MetaMorph software (Molecular Devices). Average ablation time was less than 3 seconds and time-lapse image acquisition was initiated immediately after ablation.

### Live image Acquisition

All live imaging was performed at room temperature with the following microscopes

1. Revolution WD systems (Andor Technology Ltd.) mounted on a Leica DMi8 (Leica Microsystems Inc.) with a 63x/1.4 NA objective lens under the control of MetaMorph software (Molecular devices). Images were captured using 488nm and/or 561nm lasers with a Yokogawa CSU-W1 confocal spinning disc head attached to an Andor iXon Ultra 897 EMCCD camera (Andor Technology Ltd.). Time-lapse images were acquired with 17-20 μm stacks/0.25 μm steps. Images were acquired every 30 sec for 15 min and then every 60 sec for 15 min.
2. UltraVIEW VoX Confocal Imaging System (Perkin Elmer, Waltham, MA, USA) mounted on a Nikon Eclipse Ti (Nikon Instruments, Melville NY,USA) with a 60x/1.4 NA objective lens under the control of Volocity software(v.5.3.0, Perkin-Elmer). Images were captured using 488nm and/or 561nm lasers with a Yokogawa CSU-X1 confocal spinning disc head attached to a Hamamatsu C9100-13 EMCCD camera (Perkin-Elmer, Waltham,MA,USA). Time-lapse images were acquired with 17-20 μm stacks/0.25 μm steps. Images were acquired every 30 sec for 15 min and then every 60 sec for 15 min.
3. Dragonfly 200 High-speed Confocal Imaging Platform (Andor Technology Ltd.) mounted on a Leica DMi8 (Leica Microsystems Inc.) with a 100x/1.4 NA objective lens under the control of Fusion version 2.3.0.36 (Oxford Instruments). Images were captured using 488nm and/or 561nm lasers with an Andor iXon Life 888 EMCCD camera (Andor Technology Ltd.) with the 2x mag changer. Time-lapse images were acquired with 10 μm stacks/0.25 μm steps every 1min for 30min and deconvolved using ClearView (Oxford Instruments) deconvolution.

### Live photoconversion acquisition

Photoconvertion for sM3MCA was performed using a Mosaic (Andor Technology Ltd.) on a Revolution WD system. Selected regions of interest (ROIs) were activated by a 405 nm laser for 80-90 msec.

### Live super resolution image acquisition

Super resolution imaging was performed using a VT-iSIM (VisiTech International Ltd.) mounted on a Leica DMi8 (Leica Microsystems Inc.) with a 100x/1.4 NA objective lens under the control of MetaMorph software (Molecular Devices). Images were acquired using a 488 nm laser using Orca-Fusion C14440-20UP camera (Hamamatsu Photonics). Time-lapse images were acquired with 10 μm stacks/0.25 μm steps every 15 sec for 15 min or 15sec for 7.5min followed by every 1min for 27min and deconvolved using the Fiji plug-in Microvolution (Microvolution LLC).

### Image processing, analysis, and quanitification

Image processing was performed using FIJI software (Schindelin et al., 2012). In all images, the top side is anterior and the bottom side is posterior of embryos. Kymographs were generated using the crop feature to select ROIs of 5.3 × 94.9 μm. Wound area was manually measured using Fiji and the values were imported into Prism 8.2.1 (GraphPad Software Inc.) to construct corresponding graphs.

For fluorescent lineplots, the mean fluorescence profile intensities were calculated from 51 equally spaced radial profiles anchored at the center of the wound, swept from 0° to 180°. Radial profiles of 301-pixel diameter were used. For Video S4, a diameter of 600 pixels was used to generate the line plots. Fluorescence intensity profiles were calculated and averaged using an in house code (available at: https://github.com/FredHutch/wound_radial_lineplot) using MATLAB R2020b (MathWorks), then plotted as previously described using R (Nakamura et al., 2017). For dynamic lineplots, we generated fluorescent profile plots from each timepoint and then concatenated them. The lines represent the averaged fluorescent intensity and gray area is the 95% confidence interval. Line profiles from the left to right correspond to the top to bottom of the images unless otherwise noted.

Figures were assembled in Canvas Draw 6 for Mac (Canvas GFX, Inc.).

### Actin Circle Quantification

Actin circles were counted using 300×300 pixel areas (380µm^2^) of the deconvolved single z-slice actin cortex. Two non-overlapping cropped areas were used per embryo.

### Texture Analysis

Actin architecture was assessed using cropped sections of deconvolved single z-slice images 0.25-0.5µm from the surface of the cortex of the actin ring at 70% wound closure. Gray-level co-occurrence matrix of eight 50×50 pixel cropped areas of the wound edge were used per embryo to compute four texture-related statistical properties: variance/contrast (the intensity between a pixel and its neighbor), correlation (linear dependency), uniformity (proficient order in a whole image), and homogeneity (the tightness of distribution) (Haralick et al., 1973), similar to previously described (Nakamura et al., 2020). Textural features were extracted using MATLAB R2020b (MathWorks) and were graphed on radarcharts and dotplots using R.

### Filament Orientation

Filament orientation was determined using a method similar to that described previously (Li and Munro, 2021) using Matlab 2020b (MathWorks). First, 400×400 pixel deconvolved single z-slice images of the actin ring at 70% wound closure was selected to determine the orienation of actin filaments. The images were convolved using the sobel operator and the gradient intensity and gradient direction from the DV axis was retrieved. The orthogonal of the intensity gradient direction is the filament orientation. To focus on the filaments at the edge of the wound we limited the analysis to the pixels greater than 150 pixels and less than 200 pixels from the center of the wound. Additionally, we placed a threshold based on the magnitude of the intensity gradient to remove background noise. The actin filament distribution was plotted on a radial histogram for visualization. Additionally, the orientation bias was calculated as a ratio of the density of filaments with orientations between 80°-90° and 0°-10°. This was calculated and graphed in R.

### Quantification of mRNA levels in RNAi mutants

RNAi knockdown efficiency was quantified using either qPCR or western blots. To harvest total RNA for qPCR, 100–150 embryos were collected after a 30 min incubation at 25°C, treated with TRIzol (Invitrogen/Thermo Fisher Scientific) and then with DNase I (Sigma). 1 μg of total RNA was reverse transcribed using the iScript gDNA Clear cDNA Synthesis Kit (Bio-Rad). RT-PCR was performed using the iTaq Universal SYBR Green Supermix (Bio-Rad) and primers obtained from the Fly Primer Bank listed in Table S1. Each gene in question was derived from two individual parent sets and run in two technical replicates on the CFX96 Real Time PCR Detection System (Bio-Rad) for a total of four samples per gene. RpL32 (RP-49) was used as reference genes and the knockdown efficiency (%) was obtained using the ΔΔCq calculation method compared to the control (Vermilion only) (Fig. S1F). Dia RNAi knockdown efficiency was previously shown to be 84% via western blot (Abreu-Blanco et al., 2014). Wash RNAi1 and RNAi2 knockdown efficiency was previously identified to be 99% and 96%, respectively, via western blot (Verboon et al., 2015a; Verboon et al., 2015b).

### Statistical analysis

All statistical analysis was done using Prism 8.2.1 (GraphPad, San Diego, CA). Statistical significance was calculated using a Student’ s t-test with p<0.05 considered significant.

## Acknowledgements

We thank Jeffrey Verboon for advice and comments on the manuscript, and Tony Cooke for his microscope expertise. We thank T. Allen for technical help at the start of the project, and FlyBase, the Bloomington Stock Center, the Harvard Transgenic RNAi Project, and the Vienna Drosophila Stock Center for information, DNAs, flies, and other reagents used in this study.

## Author Contributions

J. Hui, M. Nakamura, and S.M. Parkhurst contributed to the design of the experiments, performed experiments, and analyzed data. J. Hui and J.Dubrulle performed morphometrics analyses. J. Hui and S.M. Parkhurst wrote the manuscript with input from all authors.

## SUPPLEMENTARY FIGURE LEGENDS

**Figure S1.**
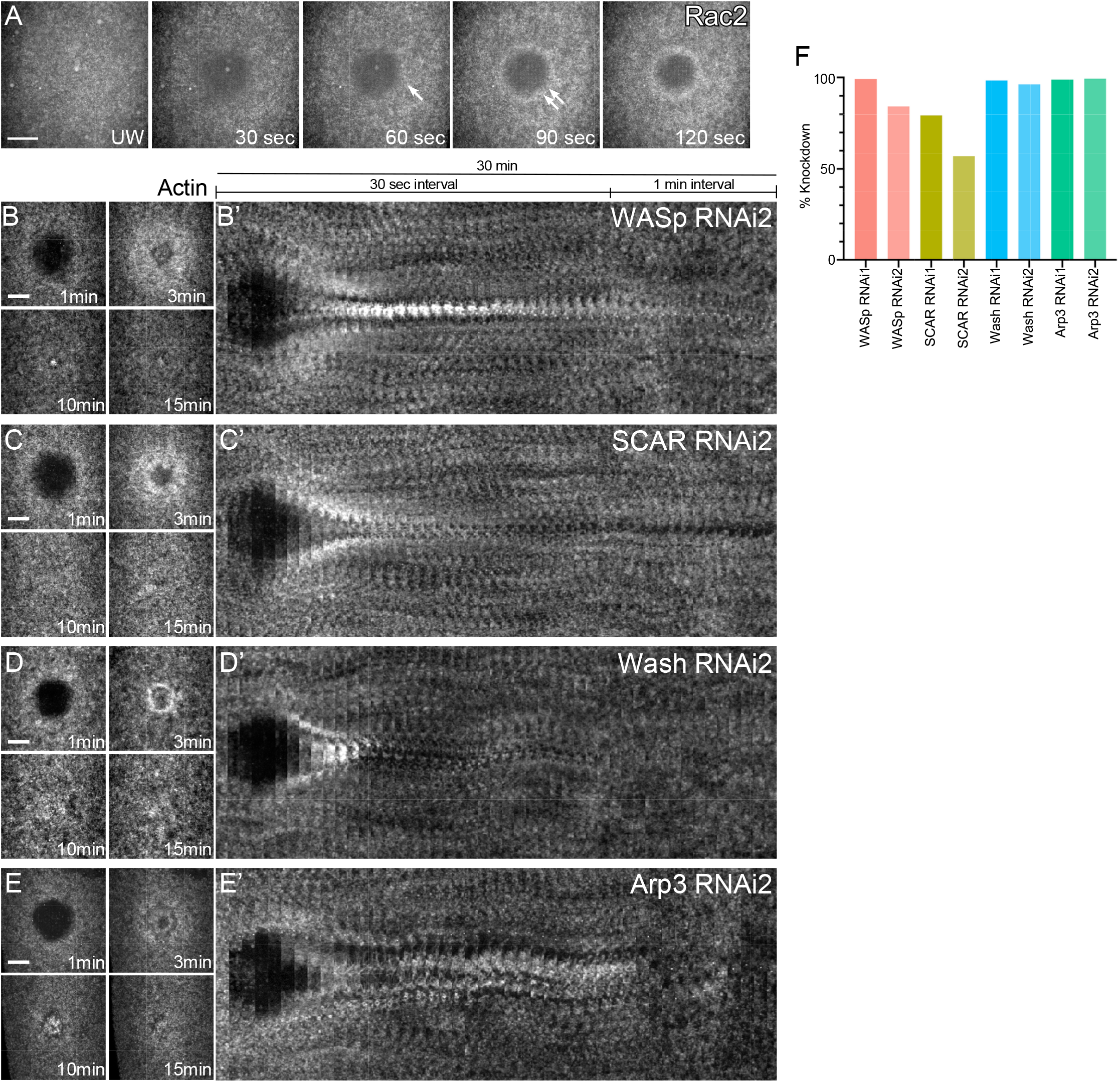
Branched actin is required for cellular wound repair and knockdown of each WAS family member exhibits non-redundant phenotypes. **(A)** XY max projections of GFP-Rac2 recruitment to the wound edge at the indicated times post wounding. **(B-E’)** Actin dynamics (sGMCA) during cell wound repair: WASp RNAi #2 (B), SCAR RNAi #2 (C), Wash RNAi #2(D), and Arp3 RNAi #2 (E). (B’ -E’) XY kymograph across the wound area depicted in B-E, respectively. Scale bar: 20µm. **(F)** RNAi knockdown efficiency quantified using qPCR.

## SUPPLEMENTARY VIDEO LEGENDS

Video 1. **Actin recruitment to wounds is defective in Rac inhibited embryos**.

(A-B) Time-lapse confocal xy images and fluorescence intensity (arbitrary units) profiles across the wound area from *Drosophila* NC4-6 staged embryos expressing an actin marker (sGMCA): control (buffer only; A) and NSC 23766 injected (B) embryos. Time post-wounding is indicated. UW: unwounded.

Video 2. **WASp, SCAR, and Wash exhibit distinct localization patterns in cell wound repair**.

(A-C) Time-lapse confocal xy images and fluorescence intensity (arbitrary units) profiles across the wound area from *Drosophila* NC4-6 staged embryos co-expressing an actin reporter (sStMCA, magenta) along with GFP-WASp (green) (A), GFP-SCAR (green) (B), or GFP-Wash (green) (C). Time post-wounding is indicated. UW: unwounded.

Video 3. **WASp, SCAR, and Wash mutants exhibit distinct phenotypes in cell wound repair**.

(A-I) Time-lapse confocal xy images and fluorescence intensity (arbitrary units) profiles across the wound area from *Drosophila* NC4-6 staged embryos expressing an actin marker (sGMCA): control (Vermilion RNAi; A), WASp RNAi 1 (B), WASp RNAi 2 (C), SCAR RNAi 1 (D), SCAR RNAi 2 (E), Wash RNAi 1 (F), Wash RNAi 2 (G), Arp3 RNAi 1 (H), and Arp3 RNAi 2 (I). Time post-wounding is indicated. UW: unwounded.

Video 4. **In the absence of branched actin and myosin, linear actin filaments swirl to close the wound**.

Time-lapse confocal xy images and fluorescence intensity (arbitrary units) profiles across the wound area from *Drosophila* NC4-6 staged Arp3 RNAi knockdown embryos treated with the Y27632 inhibitor expressing an actin marker (sGMCA). Time post-wounding is indicated. UW: unwounded.

**Table S1.**
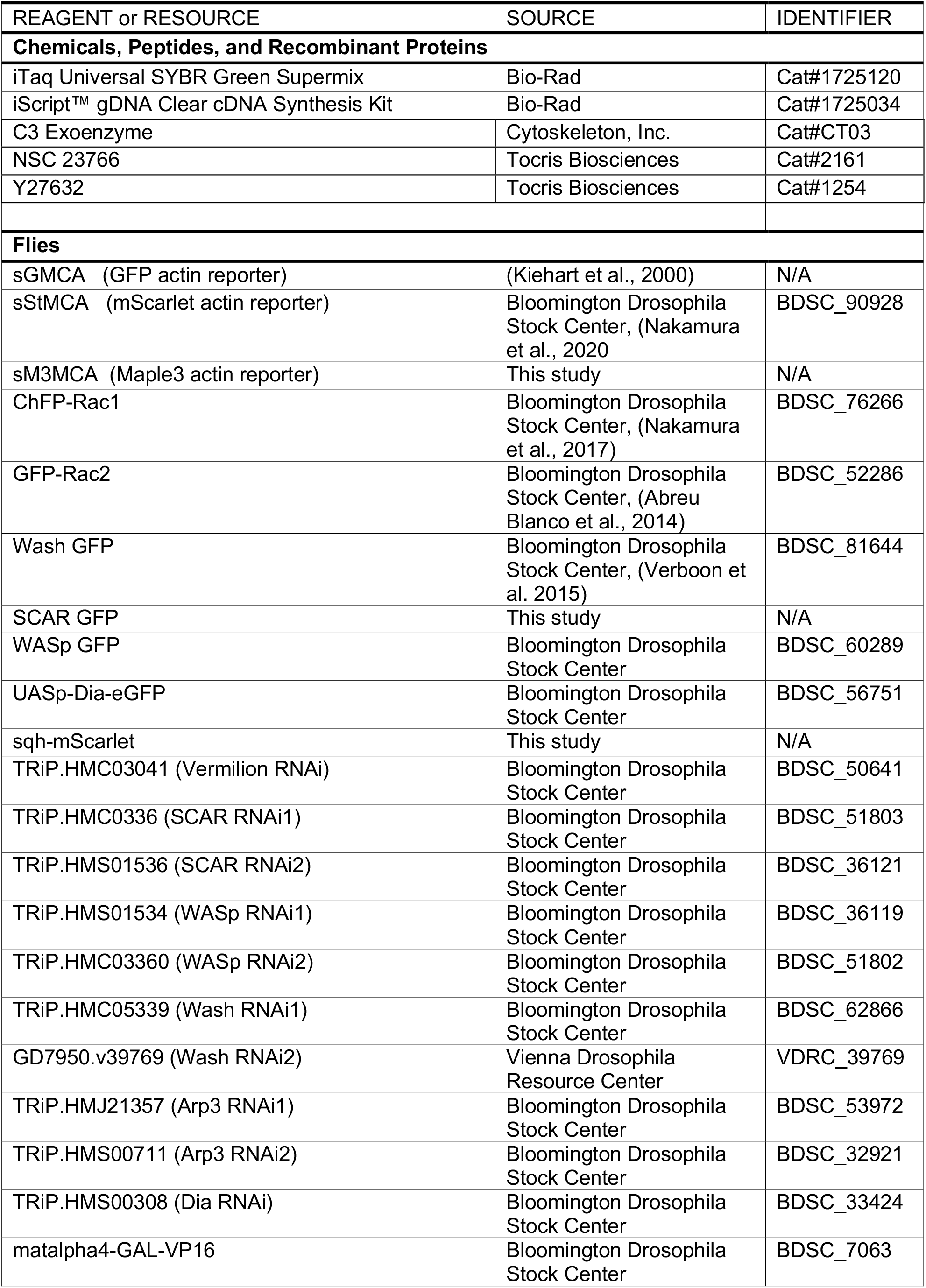
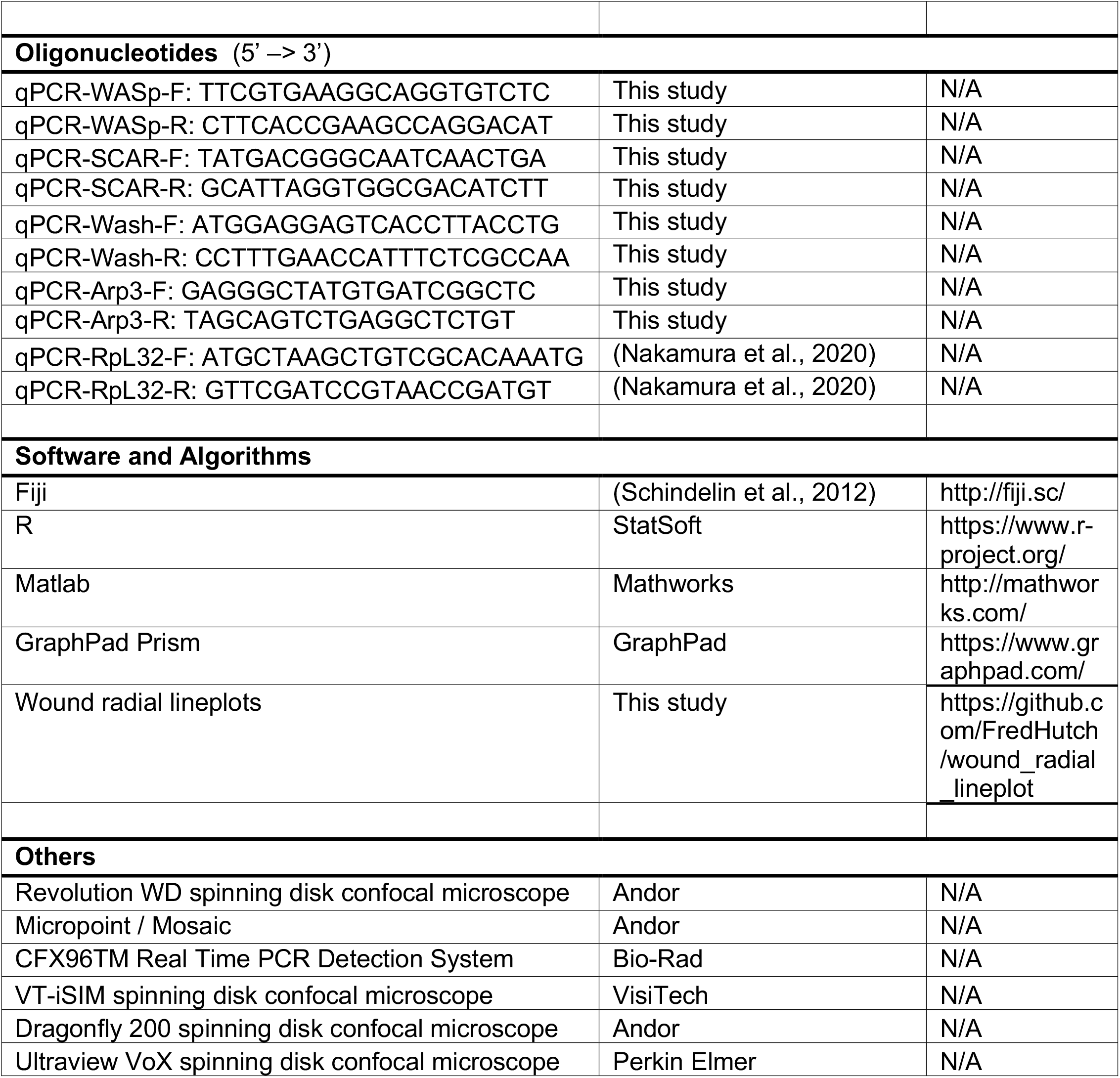
Flies and reagents used in this study

